# Understanding double descent through the lens of principal component regression

**DOI:** 10.1101/2021.04.26.441538

**Authors:** Christine H. Lind, Angela J. Yu

## Abstract

A number of recent papers have studied the double-descent phenomenon: as the number of parameters in a supervised learning model increasingly exceeds that of data points (“second-descent”), the empirical risk curve has been observed to not overfit, instead decreasing monotonically, sometimes to a level even better than the best “first-descent” model (using a subset of features not exceeding the number of data points). Understanding exactly when this happens and why it happens is an important theoretical problem. Focusing on the over-parameterized linear regression setting, a commonly chosen case study in the double-descent literature, we present two theoretical results: 1) final second-descent (regression using all of the predictor variables) and principal component (PC) regression without dimensionality reduction are equivalent; 2) the PCR risk curve can be expected to lower bound not only all linearly transformed first-descent models, but also all linearly transformed second-descent models (including the elimination of features as a special case); 3) if the smallest singular value of the design matrix is “large enough” (we will define mathematically), final second-descent can be expected to outperform any first-descent or second-descent model. These insights have important ramifications for a type of semi-supervised learning problem, a scenario which can explain why a face representation trained on unlabeled faces from one race would be better for later supervised-learning tasks on the same race of faces than for faces from another race – this can both provide a scientific explanation for the other-race effect seen in humans and give hints for how to mitigate similar issues in the domain of ethical AI.

## 1 Introduction

A number of recent studies (Belkin et al., 2018, 2019, 2020; Bartlett et al., 2020; Hastie et al., 2019) have investigated the double descent phenomenon in supervised learning: in the classical “first-descent” regime, where the number of model parameters is fewer than the number of data points, the empirical risk curve is U-shaped and thought to reflect a bias-variance trade-off (Belkin et al., 2019); however, as the number of features is increased beyond the *interpolation threshold*, along with some form of norm-minimization (usually L2 norm) to uniquely determine the solution, an over-parameterized model has sometimes been observed to monotonically decrease in empirical risk (“second descent”). It has been suggested that double descent might explain why massive neural network models have been so successful in modern machine learning (Belkin et al., 2019). Understanding when and how double descent happens is an important theoretical problems, with broad practical implications.

While the double descent phenomenon has been observed empirically in a number of settings (Belkin et al., 2019, 2018), existing theoretical analyses have primarily focused on the special case of an over-parameterized linear regression model (Belkin et al., 2020; Bartlett et al., 2020; Hastie et al., 2019) due to its analytical tractability. In this paper, we also focus on the over-parameterized linear regression problem. We approach the problem by relating it to the classical principal component regression (PCR) algorithm (Hotelling, 1957; Kendall, 1957). PCR is a well-studied “first-descent” regression technique in which the predictor variables are linearly transformed into the principal component (PC) coordinate system, and then the top *p* PCs are retained for regression (*p* ≤ *m*, where *m* is the number of data points). As we will show, understanding the theoretical relationship between PCR and double-descent will reveal new insight into the nature of the second-descent solutions, as well as the conditions under which the final second-descent can be guaranteed (in expectation) to outperform any first-descent or second-descent models that allow a linear transformation of the predictor variables.

Theoretically, the works most closely related to ours are those of Xu and Hsu (2019) and Hastie et al. (2019). At first glance, our work might seem the most similar to that of Xu and Hsu (2019) as both works attempt to gain insight into the double descent phenomenon from the perspective of PCR. However Xu and Hsu (2019) analyze an “oracle” estimator, based on the true known covariance matrix, in an (asymptotic) PCR setting. In contrast, this paper uses standard PCR, based on the sample covariance matrix, which turns out to have a much more exact and thus helpful relationship to double descent. Our setting is perhaps most similar to the equi-distributed anisotropic case of Hastie et al. (2019). However, while they claim that the behavior in the anisotropic case is *qualitatively* similar to the isotropic case, they also suggest that there is no special relation between ***β*** and **Σ** in the anisotropic case. We will show that, in PCR, such a quantitative relationship does exist. We choose to use this relationship to 1) relate the final second descent solution to the first descent PCR solutions; 2) develop conditions under which the final second descent solution is *guaranteed* to outperform any other solution (independently of feature representation) in expectation.

While the above-mentioned papers focused on developing theoretical conditions for double descent to perform well in different settings, we tackle the deeper problem of the *nature* of the second-descent solutions in relation to first-descent solutions – understanding this relationship will in turn allow us to define a novel set of conditions that can characterize guarantee final double-descent performance. In particular, if the smallest singular value of the design matrix is “large enough” (we will define this mathematically), final second-descent can be expected to outperform any first-descent or second-descent model (that allows a linear transformation of the design matrix, including the special case of feature restriction).

Our main contributions are as follows:

- We show that the final second descent solution in the linear regression setting is exactly identical to PCR using all PCs (no dimensionality reduction).
- We extend known results in PCR to the second descent to 1) show that PCR performance upper bounds linear regression performance in both descents, and 2) find conditions under which the second descent is guaranteed to outperform first descent independent of feature representation (in linear regression, as well as in PCR).
- We show how performance degrades as we move away from the optimal PCR representation, and how this might contribute to racial bias in deep convolutional neural networks (DCNNs), as well as the other race effect and social trait judgements in humans.

## 2 Preliminaries

### Notation

For a vector 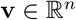 let 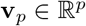 denote the sub-vector of the first *p* entries of **v**, and let 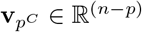 denote the sub-vector of the last *n* – *p* entries. Similarly for a matrix 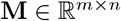, let 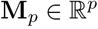 denote the sub-matrix of the first *p* columns of **M**, and let 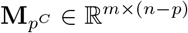 denote the sub-matrix of the last (*n* – *p*) columns.

### 2.1 Linear Regression

We consider a linear regression problem with the standard linear-Gaussian setup. We assume the data consist of *m* observations (x_1_, *y*_1_), …, (x*_m_, y_m_*) from 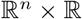 with a linear relationship

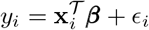

where *ϵ*_1_, … *ϵ_m_* are i.i.d 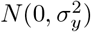 noise variables and 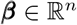 is the true coefficient vector.

Let 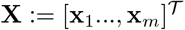 be the *m* × *n* design matrix and let 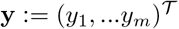 be the m-dimensional response vector. We assume without a loss of generality that the rank of **X** is min(*m, n*), and that **y** is centered.

The linear regression estimator 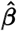 of ***β*** is then

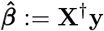

where the symbol ‡ denotes the Moore-Penrose pseudoinverse. Given new test data 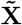, their responses can be estimated as 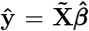. In the first descent (*n* ≤ *m*), 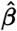 is the least-square estimator; in the second descent (*n* > *m*), this estimator has the minimal L2 norm among all interpolating solutions (training error = 0).

### 2.2 Principal Component Regression

Principal component regression (PCR) (Hotelling, 1957; Kendall, 1957) is a regression technique based on principal component analysis (PCA) (F.R.S., 1901; Hotelling, 1933, 1936), whereby the projection of the design matrix **X** along the PCs are used as the predictor variables in the regression analysis. One can choose to either use all of the PCs or a subset of PCs, usually the PCs with the *p* largest projected sample variance (equivalently, the *p* largest eigenvectors of the sample covariance matrix).

Recall that the design matrix **X** can be decomposed in terms of its singular value decomposition (SVD) as

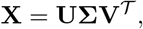

where **U** and **V** are unitary (orthogonal if real-valued) matrices whose columns consisting of left and right singular vectors, respectively, and **Σ** is a diagonal matrix of singular values. Note that the first *m* columns of **V** are also the principal components (PCs) of **X**. The projection of **X** onto the PCs is thus **Z** := **XV***_m_*. Note that, since **V***_m^C^_* is in the null space of **X** (i.e. **Σ***_m^C^_* = **0**), it follows that **Z** = **XV***_m_* = **XV**.

PCR assumes the first *p* ≤ *m* PC dimensions are retained for regression. The PCR estimator 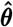 of ***θ*** is thus

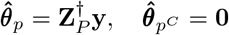

It follows from the definition of **Z** that an alternative expression for 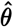, in terms of 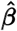, is just 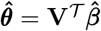.

### 2.3 Relevant PCR Properties

We state below several useful properties of PCR that will help us gain insight into double descent.

#### PCR1. Optimality of PCR in the first descent

Suppose we wanted to find the best first-descent linear regression model for (**X**, **y**) in the sense of minimizing expected MSE on test data, 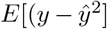, by allowing a linear transformation of **X** and restricting to *k* ≤ *m* features. Park (1981) showed that if only *k* features are allowed in the linear regression, then the best such linear transformation is PC projection using the *k* largest PCs, i.e. Λ*X* = **XV***_k_*, where **V***_k_* consists of the first *k* right singular vectors.

#### PCR2. Optimizing number of PC features in PCR

Furthermore, if one wants to optimize for *k* so as to minimize test MSE, Park (1981) showed that, for any *p* ≤ *m* and any matrix 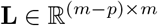 of rank (*m* – *p*),

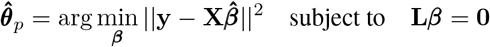

the *p*-th PC should be included if and only if *σ_p_* (the *p* – *th* singular value) satisfies the following threshold:

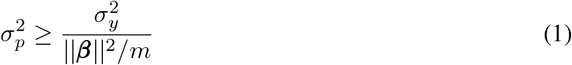

Here, *σ_y_* is the noise in **y**, ‖***β***‖ is the norm of the true parameter vector ***β***, and *m* is the number of entries in ***β***. The entries of ***β*** are assumed to have a uniform distribution over all directions. See Park (1981) for a detailed proof.

Note that the relationship in Eqn. 1 can be viewed as a form of “signal-to-noise” ratio of the *p*-th dimension (SNR*_p_*), whereby the noise is 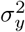 and the signal in dimension *p* is 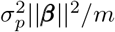. Note that, in the isotropic case (**Σ** = **I**), the relationship in Eqn. 3 simplifies to the SNR relationship developed by Hastie et al. (2019) (up to normalization constant *n*).

Together, PCR1 and PCR2 imply that the process of feature selection in PCR can be done sequentially and greedily: proceed from the the largest to the smallest PC, stop adding features as soon as inequality 1 is not satisfied.

#### PCR3. Nested nature of PCR models

Let 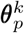 be the first *k* weights of the parameter vector obtained using the first *p* PCs, let 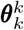 be the first *k* weights of the parameter vector obtained using the first *k* PCs with *k* ≤ *p*. Then 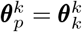.

This follows from **Z** = **UΣ** (or equivalently **V** = **I**).

Let **Z** = **UΣ** be the SVD decomposition of 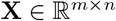, and let 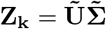 be the SVD decomposition of **X** using only the first *k* ≤ *n* features. Then 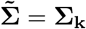 and 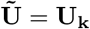.

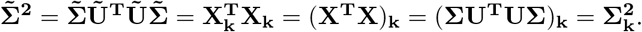

Since 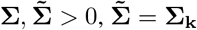.

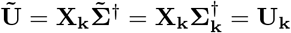

Then 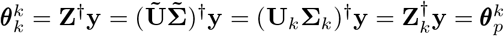

## 3 Insight into Double Descent from PCR

We will leverage the results summarized in the previous section to gain some insight into double descent.

### Lemma 1. Equivalence of PCR and final second-descent

Let 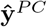 denote predictions made with all *m* PCs and let 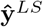 denote predictions made with all *n* original features. Then 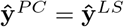. This follows directly from the orthogonality of **V** and can be shown using basic linear algebra as follows.

Let 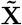 be test data in the original representation. Then the test data in the PC representation is 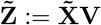, and

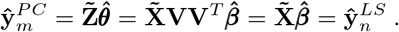

In other words, predictions using the original feature representation and the PC representation are identical. Because the choice of test data 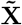 is arbitrary, it follows that the regression solutions are exactly the same between final second-descent and PCR using all PCs. Compared to PCR including all PCs, the final second descent solution is just a change of coordinate system and addition of trivial dimensions.

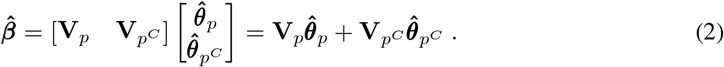

This follows from the definition of 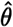, as well as PCR3.

While final second-descent might appear to have many more feature dimensions than PCR, in fact the least-L2-norm solution estimates the slope to be 0 in all dimensions orthogonal to the span of the training data **X**.

### Lemma 2. PCR lower bounds second-descent

Following from Lemma 1, if some of the original features are eliminated so that only *p* (*m* < *p* < *n*) features are retained for regression, then the *p*-feature regression solution is equivalent to doing regression on PCs not computed from the full *n* features, but from only the *p* features, which can have rank at most *m*. In other words, every second-descent regression solution is exactly equivalent to some first-descent regression solution (obtained by linearly transforming the original feature representation). Suppose the rank of **X***_p_* is *h*, then the expected MSE for this algorithm can be expected to be lower-bounded by the MSE of he *h*-PC PCR solution (due to PCR1).

### Theorem 1.

The MSE of the best PCR model lower bounds the entire double-descent MSE curve. This follows directly from PCR1 and Lemma 2.

### Theorem 2.

Assume the entries of ***β*** to have a uniform distribution over all directions. If the smallest singular value *σ_m_* of the representation satisfies the following threshold:

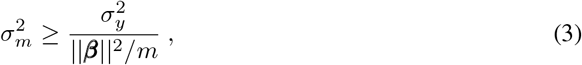

where *σ_y_* is the noise in **y** and ‖***β***‖ is the norm of the true parameter vector ***β***, then the final second-descent solution is guaranteed (in expectation) to outperform any other first- or second-descent solution obtained by linearly transforming the original feature representation of the predictor variables **X**).

Theorem 2 follows from PCR2 and Theorem 1.

See the Appendix for numerical simulations that illustrate the theorems.

### 3.1 Simulations

Here, we observe what happens as the feature representation is rotated away from the optimal PC representation. We expect regression performance on test data to degrade gradually.

Figure 1 shows how regression performance degrades as the feature representation is slowly rotated away from the PCR representation by 1) adding Gaussian noise 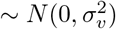 to **V** and then orthogonalizing (Figure 1a) and 2) increasing data points added to the PCA transformation, so that the PCs correspond to those of the larger super set and look increasingly less like the PCs of only the training data (Figure 1b) – it is done by performing PCA on the extended design matrices in 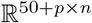 to yield a 50 + *p* dimensional representation. Note that the performance degradation affects both first and second descent, but are invariant with respect to the manipulations for PCR including all PCs and final second-descent.

**Figure 1:**
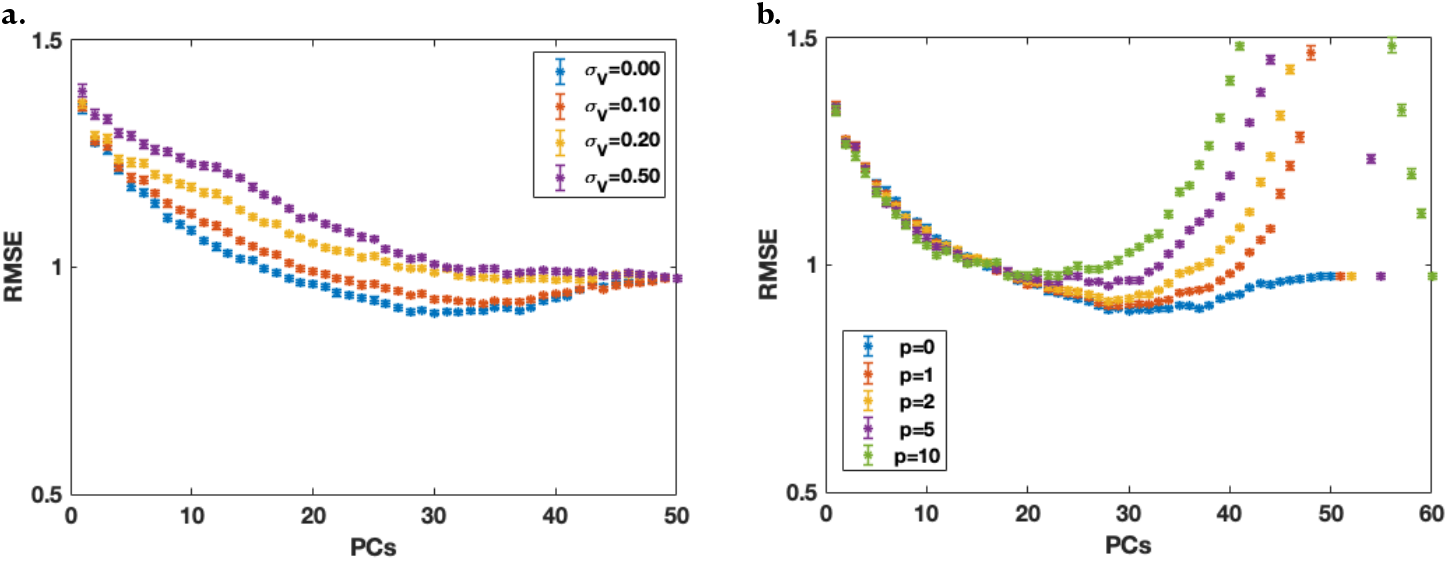
a,b) Root mean square error (RMSE) vs. number of parameters (PCs) in two modified PCR settings on test data (10 datapoints) averaged over 25 different design matrices 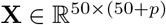. In both a) and b) performance degrades from the PCR baseline (*σ_v_* = 0 and *p* = 0). Note that in both a) and b) the baseline is identical to the second case in Figure 4a) (SNR*_m_* = 0.5) in the Appendix. Errorbars are standard error of the mean (sem).

We also observe that, in the overparameterized regime, as the PCs (**V**_*m*_) are rotated away from the PCR representation (**V**_*m*_ = **I**), the null space vectors (**V**_*m^C^*_) are rotated toward **I** (Figure 2). This is not unexpected, as **V**_*m*_ and **V**_*m^C^*_ are related to one another by 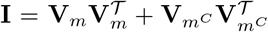. As such, by necessity, as **V**_*m*_ becomes less orthogonal, 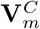 must become more orthogonal. It follows then, from the nested nature of PCR models as well as the equivalence of final second descent solutions, that **V**_*m*_ rotates 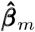 away from 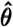, while 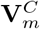 rotates 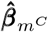 towards 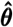. See the Appendix for a simulation.

**Figure 2:**
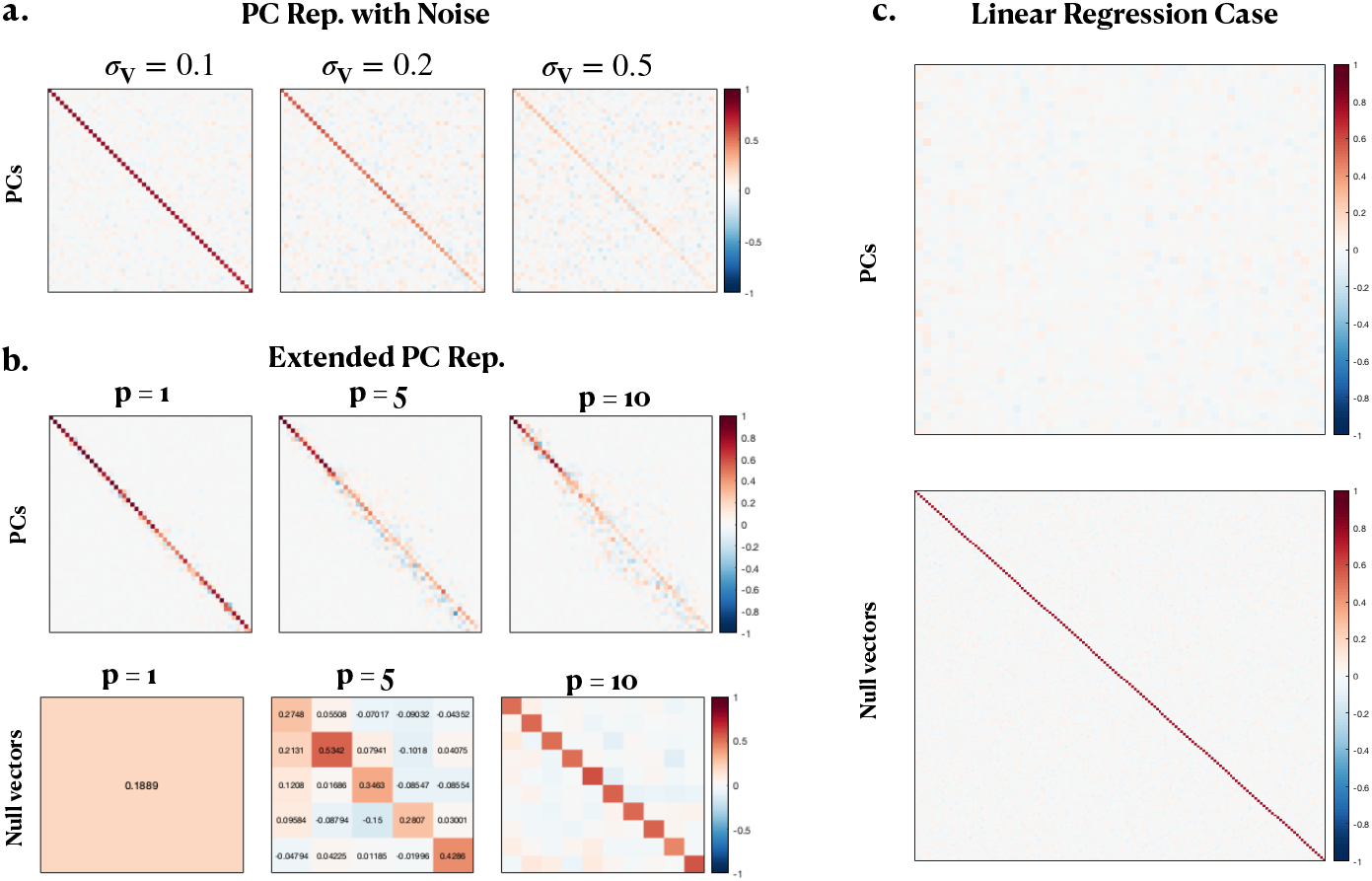
a, b) **V** resulting from the operations corresponding to Figure 1a) and 1b) respectively. c) **V** corresponding to SNR*_m_* = 0.5 in Figure 4b) in the Appendix. a,b) **V**_*m*×*m*_ loses its structure uniformly across dimensions (a) and mostly distorts the later PCs (later diagonal elements of **V**_*m*×*m*_) (b, first row). In the case of increased overparameterization, we observe that **V**_*m^C^* × *m^C^*_ (the matrix of nullspace vectors) recovers a nearly diagonal structure (b, second row). c) In the case of linear regression (performance shown in the Appendix), we observe that **V**_*m*×*m*_ has lost its diagonal structure (nearly) entirely (first row), implying that the first descent feature representation has been rotated (nearly) entirely away from the optimal PC representation, while **V**_*m^C^* × *m^C^*_ is very nearly diagonal (second row).

## 4 Other Race Effect and Semi-Supervised Learning

Human learning and machine learning both are known to suffer from poor generalization when the feature representation is ill-suited to a supervised learning problem due to mismatched data statistics. For example, humans suffer from the other-race effect (ORE) (Malpass and Kravitz, 1969; Valentine, 1991), whereby people are better at processing own-race faces than other-race faces (Meissner and Brigham, 2001) due to the relative lack of exposure to other-race faces during child development (Valentine, 1991). Similarly, modern machine learning algorithms can suffer from social biases due to a mismatch of data distribution in training and test data, e.g. deep convolutional neural networks (DCNN) trained on predominantly white faces are much worse at recognizing Asian faces (compared to White) and vice versa (Xiong et al., 2018; Tian et al., 2021). Our results on PCR and double-descent offer an interesting hypothesis why ORE happens in humans, and also for how to mitigate similar issues in machine learning algorithms.

Human face-representation learning can be thought of as a particular type of semi-supervised learning: representation learning using a large set of unlabeled data is followed by regression modeling using a smaller set of labeled data. Children have exposure to massive amounts of unlabeled data and relatively infrequent direct feedback, and their brains are highly plastic during development; past adolescence, brain plasticity rapidly declines, coinciding with a decreased ability to master new supervised learning tasks, such as learning a new language (Penfleld and Roberts, 1959) or processing a race of faces that one has little prior experience with (Young et al., 2012). Based on our theoretical findings, we expect that regression learning performs better if the smallest singular values of the labeled training data (predictor variables) are larger. We postulate that this is why a face representation trained on faces from one race would be better for later supervised-learning tasks on the same race of faces than for faces from another race.

Given the optimality of PCR for regression that we showed in this paper, it makes sense for a semi-supervised learning algorithm to learn a PC representation for the unlabeled data. It has been previously shown that experience-dependent (unsupervised) learning in the brain, based on known properties of neural plasticity, can naturally support online PC representation learning (Minden et al., 2018). If the statistics of the labeled data (predictors) are similar to the unlabeled data, we can expect the supervised learning component to do well; however, when the statistics of the labeled data are substantially different from the unlabeled data, as in the case for other-race faces, then we can expect worse supervised learning performance. We first elaborate on this concept mathematically, then demonstrate it in an experiment.

Let **X***_ul_* be the matrix of unlabeled faces, and let **X***_l_* be the matrix of labeled faces (predictor variables). Then:

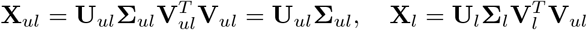

For ease of notation let 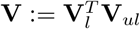, where **V***_ul_* is fixed. When the PCs of unlabeled and labeled faces are identical (**V***_ul_* = **V***_l_*, i.e. **V** = **I**), the representation is optimal also for labeled faces (**X***_l_* = **U***_l_***Σ***_l_*). However, as seen in Figures 1 and 2, when **V***_l_* is rotated away from **V***_ul_* (i.e. **V** is rotated away from **I**), then **X***_l_* is rotated away from its optimal representation.

Note that, as **V** is orthogonal, the entries **V***_ij_* are simply the dot products of the *i*-th PC for unlabeled faces and the *j*-th PC of the labeled faces. The diagonal entries of **V** thus provide an explicit measure of similarity between PCs for unlabeled and labeled sets of faces.

We postulate that ORE arises from a feature representation optimized for unlabeled faces. If this is the case, then we expect:

1. The PCs of labeled faces will be better aligned with the PCs of the unlabeled faces, when the latter are of the same race rather than of a different race.
2. The smallest singular values (sample standard deviations) of labeled faces should be larger when the feature (PC) representation is learned on unlabeled same-race faces, than when it is learned on unlabeled other-race faces.
3. Regression performance should be correspondingly better for same-race faces than other-race faces.

### Experiment

We use two different initial feature representations:

1. The activations of the last fully connected layer (FC2) from a DCNN (VGG16 (Simonyan and Zisserman, 2015)) trained on VGGFace (Parkhi et al., 2015). FC2 was chosen as it has been suggested that the activation responses of the layer prior to the final classification layer is a typical representation for each face in a DCNN (O’Toole et al., 2018; Tian et al., 2021).
2. The shape and texture features of an Active Appearance Model (AAM) (Cootes et al., 1998; Edwards et al., 1998) trained on the 10k Adult Faces database (Bainbridge et al., 2013) as well as the CFD dataset (Ma et al., 2015). AAM was chosen as it has been shown that AM cells in the macaque brain approximately encode linear combinations of the axes (Chang and Tsao, 2017) and it allows for easy visualization of its feature dimensionsGuan et al. (2018).

We then complete “unsupervised learning” by performing PCA on a set of unlabeled faces.

The actual face images we use are Asian, Black and White faces (104 total and 52 female for each race) from the Chicago Face Database (CFD) (Ma et al., 2015). We split the faces of each race into unlabeled (*m* = 75), labeled training (*m* = 25), and labeled testing (*m* = 4). We then project the labeled faces (same and other race) into this representation; a subset of these are used as training data to estimate the social trait ratings collected by Ma et al. (2015) with CFD, while the remaining labeled data are used to evaluate regression performance. We do 50 random splits and then average across all splits.

As shown in Figure 3 and the Appendix, we observe that the best test RMSE is obtained when the unlabeled faces (“representation”) are from the same race as the regression faces (VGG16: Figure 3a, AAM: appendix). This is true throughout both the first-descent and second-descent parts of the empirical RMSE curves. AAM allows us to visualize how the feature dimensions differ for the Asian, Black and White representations, through synthetic faces generated along the PC features: we see that the top PCs tend to capture variations consistent with same-race faces, and rarely with other-race faces (Figure 3b). Correspondingly, we also observe that the PCs of same-race faces are more closely aligned than different-race faces, with alignment measured as the average dot product between PCs (diagonal elements of **V**, see Figure 3c).

**Figure 3:**
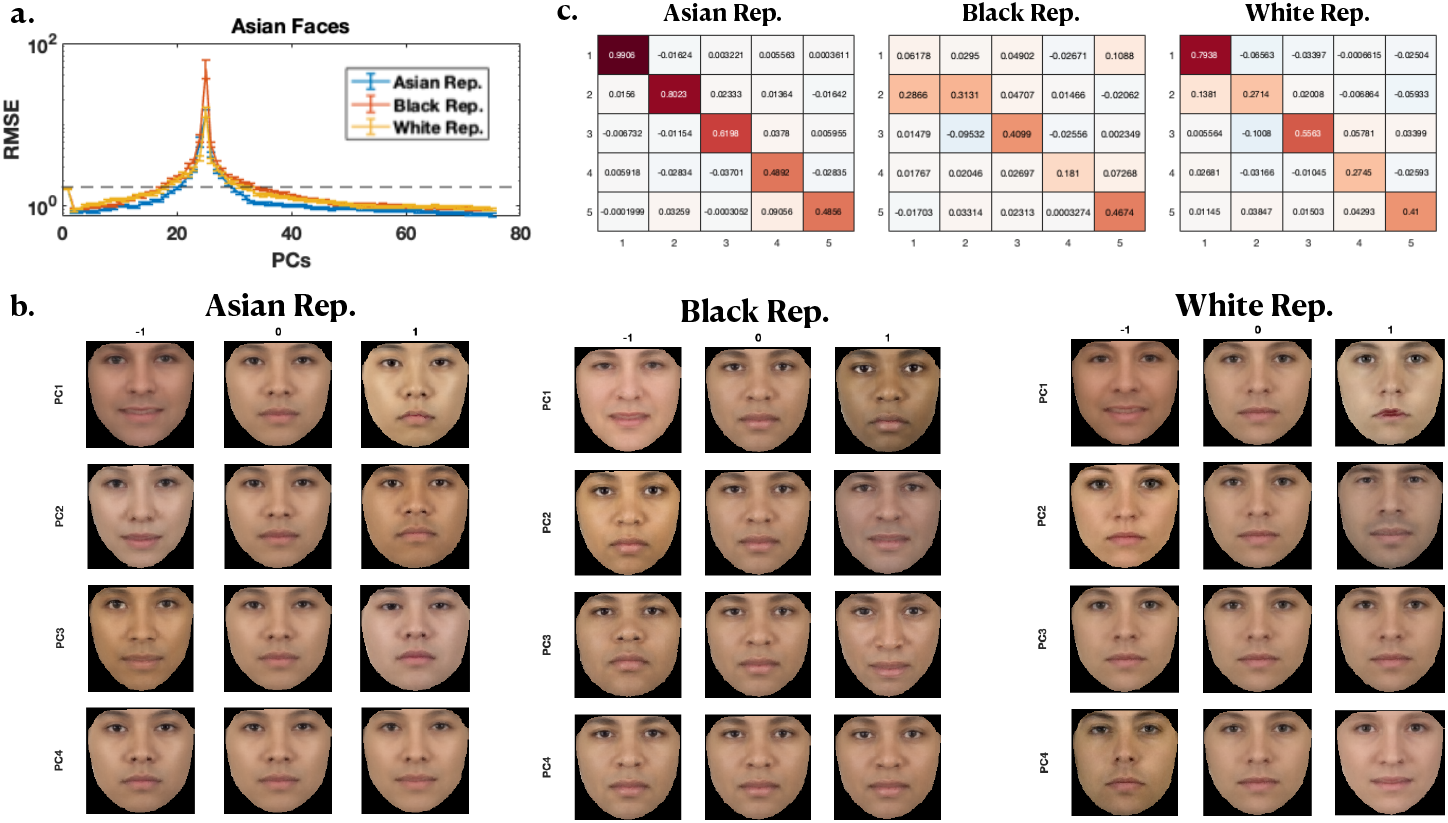
Other Race Effect. a) RMSE vs number of parameters for Asian labeled faces using an Asian (blue), Black (red) and White (yellow) unlabeled face representation for estimating the “feminine” trait from Ma et al. (2015) using the VGG16 representation. Performance is better for same race faces than other race faces, indicating that ORE is present. Black dashed line indicate standard deviation of the ratings. See the Appendix for Black and White faces. b) Average mixed-race face (averaged over all 104 Asian, Black and White faces) generated along the first four PCs of the Asian (left), Black (middle), and White (right) representations. Note that the mean faces (middle column of each representation) differ between the representations even though the same average face was used as the features differ between the representations. c) The first 5 rows and columns of **V** for Asian faces using an Asian (left column), Black (middle column) and White (left column) AAM representation. Diagonal elements of **V** (i.e. dot products (dp) of PCs) are on average larger for same race faces than other race faces, indicating that the PCs are better aligned for same race than other race faces. See the Appendix for Black and White faces.

## 5 Discussion

In this work, we identified an equivalence between overparameterized linear regression PCR. In particular, when the smallest singular value of the data matrix exceeds a particular threshold, PCR with all PCS and thus the final second-descent algorithm can be expected to outperform any first- or second-descent model (allowing arbitrary linear transformations of the predictor variables). We showed that all of the second-descent models in regression correspond to some first-descent model, whose expected tested error is lower-bounded by that of PCR – thus they are not a new fancy class of regression models that what have been studied in the classical first-descent regime. The reason why performance tends to improve in the second descent is because their equivalent first-descent algorithms become closer to exact PCR. We also gained some insight as to why PCR works so well and does not suffer from nearly as much over-fitting as other representations: (1) PCR’s solutions are nested, so that adding the smallest PCs do not change the estimated coefficients for the larger PCs and thus their overall contribution and in particular the ability to induce over-fitting is quite limited, since by definition test data magnitude and thus predictive influence can be expected to be much smaller in these dimensions; in particular, if the smallest singular value is “large enough”, even including the smallest PC features; (2) any over-parameterized transformation of the training PCs can be expected to perform worse than PCR and leader to a higher peaking of MSE at the interpolation threshold, since it is well known that the smallest singular value of the design matrix monotonically decreases toward the interpolation threshold and decreases past the interpolation threshold – until it becomes exactly identical to the smallest singular value of PCR at final second-descent.

We also applied our theoretical insights to an important type of semi-supervised learning problem, whereby representation learning using a large set of unlabeled data is followed by regression modeling using a smaller set of labeled data. This problem setting is very useful for machine learning, as massive quantities of unlabeled data is easy to obtain while labeled data is expensive and relatively scarce. It is also the setting in which the brain excels in, as children have exposure to massive amounts of unlabeled data and relatively infrequent direct feedback. Based on our theoretical findings, we expected that regression learning preforms better if the smallest singular values of the labeled training data (predictor variables) are larger. We postulated that this is why a face representation trained on faces from one race would be better for later supervised-learning tasks on the same race of faces than for faces from another race. This in turn provides a scientific explanation for the other-race effect (Malpass and Kravitz, 1969; Valentine, 1991) seen in humans and sheds light on how to mitigate similar issues in the domain of ethical AI (Bolukbasi et al., 2016; Guynn, 2015). Our empirical results bore out these hypotheses.

Although obviously restricted to the linear regression domain, this work has interesting relevance to theoretical neuroscience and broader implications for several topical areas within machine learning. While modern machine learning has achieved incredible performance in some areas (He et al., 2015; Silver et al., 2016), relative to the human brain, it still falls short on several important respects. For example, deep neural networks are extremely labeled-data intensive and computation-intensive (Krizhevsky et al., 2012; Thompson et al., 2020). In contrast, the developing brain receives a great deal of unlabeled data but only a very small amount of labeled data, i.e. putting it more in the regime of semi-supervised learning than supervised learning. The brain also completes a large variety of visual perception tasks with high accuracy and computational efficiency (Lake et al., 2017, 2019). Related to this, deep neural networks typically generalizes poorly in terms of transfer learning (Recht et al., 2018, 2019) and few-shot learning (Oreshkin et al., 2018). But the human brain appears capable of effortlessly and accurately generalizing to completely novel perceptual tasks. The current work suggests that one way to tackle this problem may be to have a massive over-parameterized feature representation that is largely learned in an unsupervised manner, and then train another decoding layer *as needed* for a novel supervised learning task (e.g. regression). The results (final second-descent) can be guaranteed to be as good as or better than any specialized model that does feature selection for each supervised learning task, based on the results we described here. Interestingly, it has been shown that across species and sensory modalities, the brain tends to massively expand the feature representation from the initial sensory receptor level (Olshausen and Field, 1997; Dasgupta et al., 2018). Previously, it was suggested that such feature expansion may serve some appealing goals within unsupervised learning, such as reducing energetic needs in the brain (sparsification) or preserving semantic similarity among sensory input. However, our result suggests that this feature expansion in the brain may instead serve as a powerful *universal* representation that can readily support future unspecified supervised learning needs with high accuracy and sample-efficiency.

This work can be developed in multiple fruitful future directions. For example, it is worthwhile to examine how the results presented here generalize to other supervised learning tasks (e.g. classification), or relate to more complex multi-level architecture (instead of only single-level linear regression). Another interesting direction is to identify more precisely the general feature representational settings under which final second-descent can be expected to out-perform best first-descent. Yet another direction is to examine how things generalize with a different norm for minimization in the second descent. For example, in theoretical neuroscience, L1 norm (sparsification) has received more attention than L2 norm minimization in overcomplete representations (Olshausen and Field, 1997); an interesting question is which first-descetn solution L1 norm minimization might be equivalent to, and, more generally, how one should choose the norm to minimize in the second descent.

## Acknowledgements

We thank Mikhail Belkin, Sanjoy Dasgupta, Alex Cloninger, Ken Kreutz-Delgado, Srinjoy Das, Marcello Mattar, and Homero Esmeraldo for inspiring and helpful discussions.

## A Appendix

An important concept introduced in the main paper is that we can define a form of signal-to-noise (SNR), 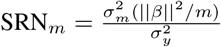, which is a minimal threshold that the smallest singular values need to exceed in order to be included in the optimal (in expectation) PCR algorithm. As SNR*_m_* decreases, we can expect that more PC features need to be dropped, and the test error rate to increase; we also expect that the empirical risk curve to become more U-shaped, as the inclusion of the smallest PCs begin to degrade performance. Figure 4a illustrates the above to be indeed true in synthetic data. Another major concept introduced in the main paper is that the PCR risk curve should lower-bound both first- and double-descent risk curve, if regression is done in the original feature space and not in the PC space. This is illustrated by simulation in Figure 4b.

**Figure 4:**
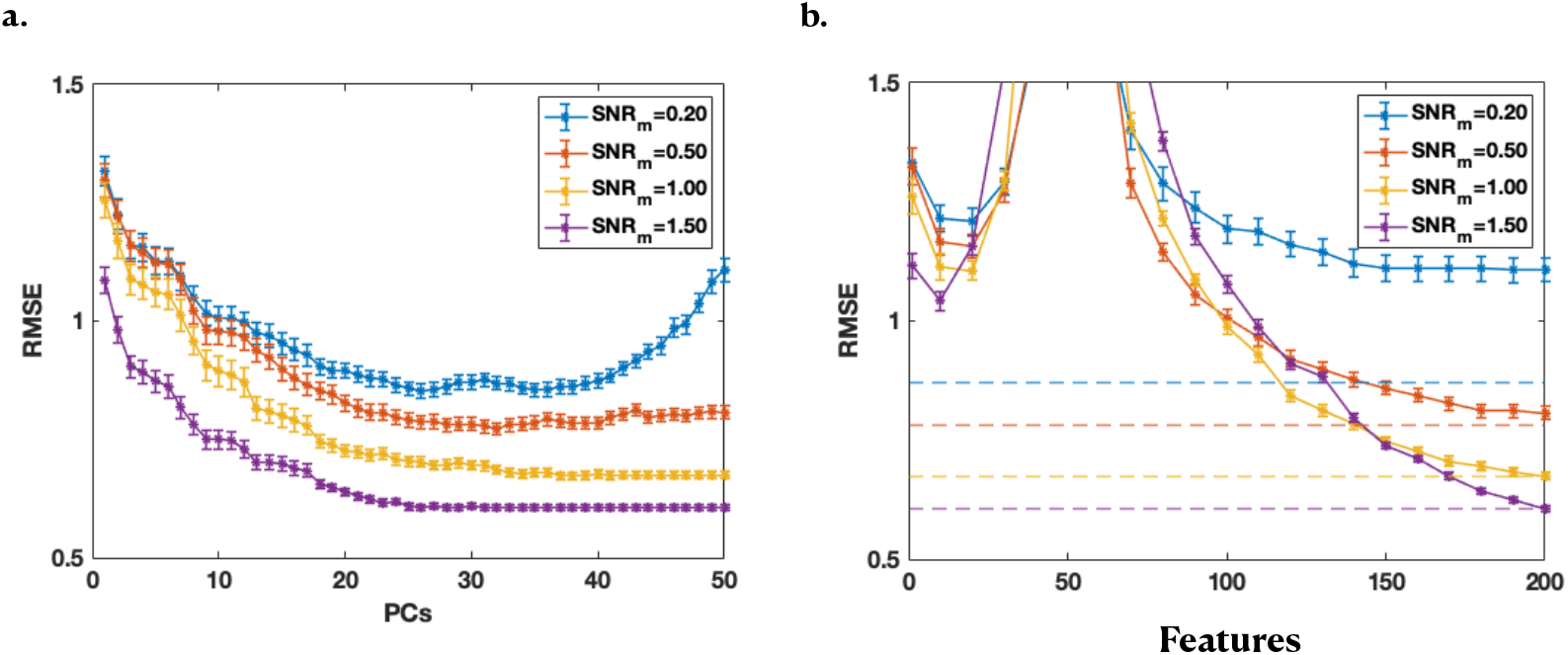
Root mean square error (RMSE) vs. number of parameters (PCs) in PCR (a) and linear regression (b) on test data (10 datapoints) averaged over 25 different design matrices 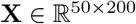. Each design matrix was generated as **X** = **UΣV***^T^*, with the diagonal entries of **Σ** ~ *Exp*(1) and thresholded using *σ_m_* ∈ {0.1, 0.25, 0.5, 0.75}. Dashed lines in (b) indicate the best performance achieved by PCR in (a). SNR*_m_* is defined by Equation 3 in the main text. To vary SNR‖*β*‖^2^/*m* and *σ_y_* are kept fixed (‖*β*‖^2^/*m* = 1 and *σ_y_* = 0.5), while *σ_m_* varies from 0.1 to 0.75. Errorbars are standard errors of the mean (sem). Note that for SNR*_m_* ≥ 1, the lower bound is tight. This is because optimal performance is achieved with the final second-descent solution, and this solution makes equivalent predictions in the two settings.

Figure 5 shows how solutions become less and then more similar to the final second-descent solution (in terms of the dot product between the *k* features of the *k*-th solution using only the first *k* features in regression, and the first *k* features of the full solution using all features), in the under- and overparameterized regimes, respectively. This is because the final second-descent solution is also identical to the final PCR solution (including all training features), after a redefinition of the basis. To create an overcomplete parameterized representation here, we generate additional *p* synthetic samples from the generative model for x, and then perform PCA on the superset of data points including the training data points for regression and these additional data points. Doing PCA on this superset of data points creates a feature representation that has more features than PCA on only the regression training data, thus allowing us to create the double-descent risk curve. The more extra data points are included in the superset (larger *p*), the more the feature representation (obtained by doing PCA on this expanded data set) shifts away from that of the exact PC representation of training data, though this affects the smaller PCs more, as the estimation of larger PCs are more robust due to the addition or elimination of a few data points. That is why the dot product of the first 40 or so features are still quite close to 1 in comparison to the final second-descent solution. In contrast, for the dot product of the baseline solution (light blue), the first-descent solutions are not so similar to the final second-descent solution (dot product far from 1 in the first descent), because that representation was not created by design to be similar to the training PC representation.

**Figure 5:**
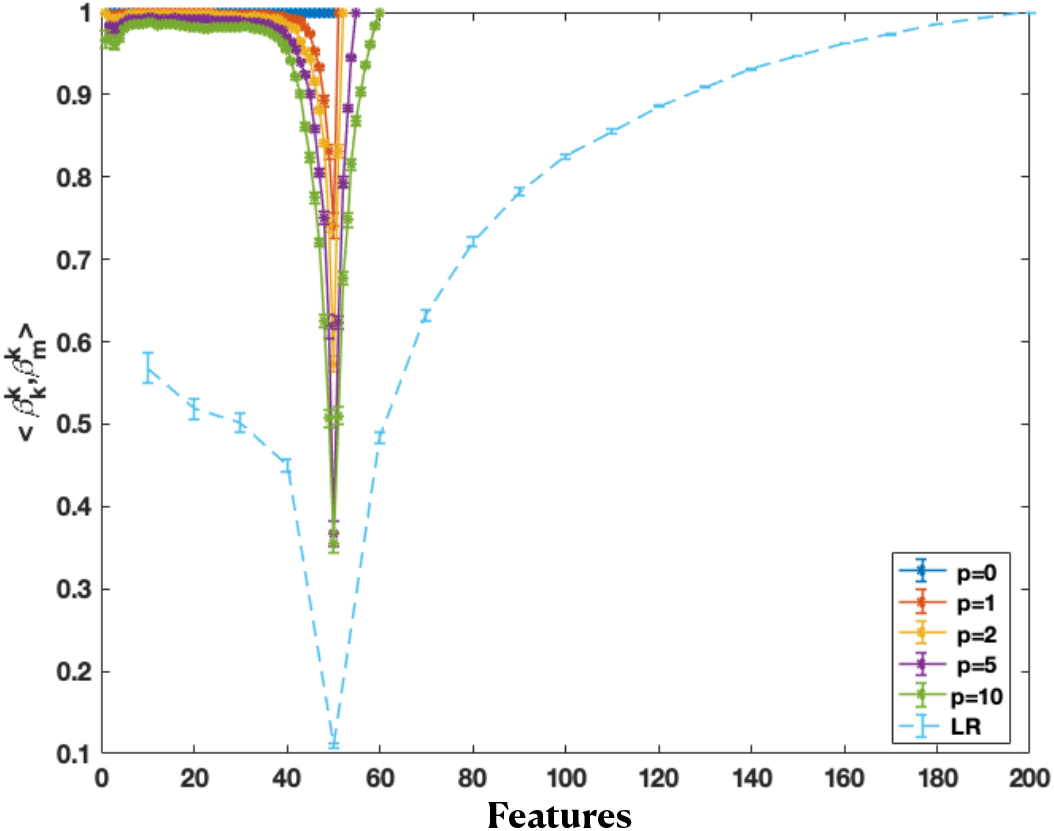
Dot product of the first *k* (PC) feature weights of the full solution and the *k*-th solution, using a (50 + *p*)-dimensional representation, as well as the linear regression case in Figure 1b) (LR). The (50 + *p*)-dimensional representation is generated by increasing the data points added to the PCA transformation (by generating more synthetic datapoints from the same distribution). This is done by performing PCA on the extended design matrices in 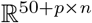 to yield a 50 + *p* dimensional representation. The dot product becomes smaller in the first descent, then larger in the second descent, indicating the solutions become less, then more similar again.

Figures 6 and 7 simulate how the other race effect (ORE) for Asian, White and Black faces can arise in VGG16 (Fig. 6) and AAM (Fig. 7). Given the original features (in either VGG16 or AAM), a linear transformation is performed on a set of “unsupervised learning” faces. Then a linear regression problem is imposed on this “unsupervised learned” representation. Figure 6a (VGG16) and Figure 7a (AAM) show that the test error is lowest in both first- and second-descent for same-race faces than for different-race faces. Figure 6b (VGG16) and Figure 7b (AAM) show that the features are closer to training PC features (*mathbfV* is more diagonal) for same-race than different-race faces.

**Figure 6:**
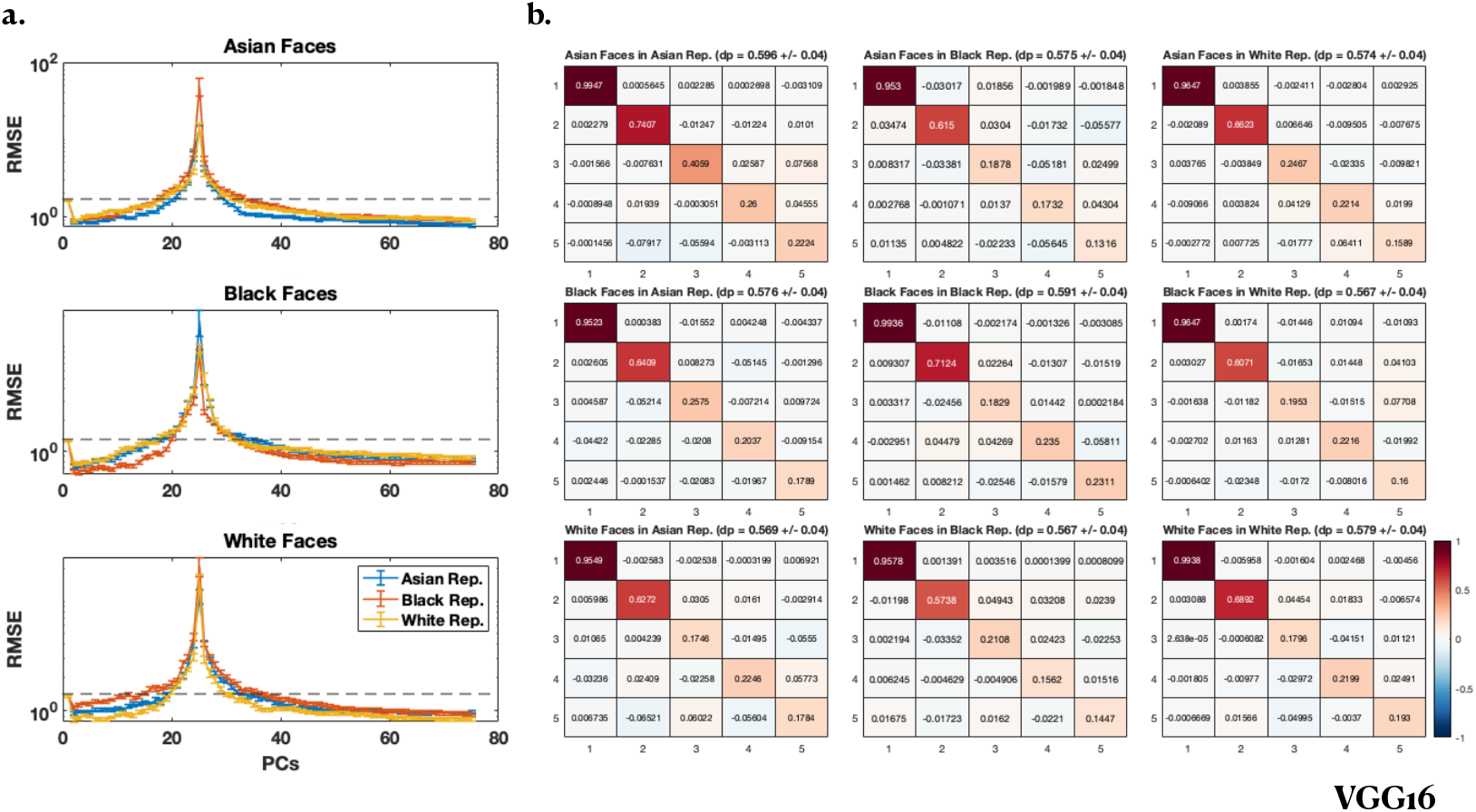
VGG16. a) RMSE vs number of parameters for Asian (top row), Black (middle row) and White (bottom row) labeled faces using an Asian (blue), Black (red) and White (yellow) unlabeled face representation for estimating the “feminine” trait from the CFD dataset (Ma et al., 2015). Performance is better for same race faces than other race faces, indicating that ORE is present in all cases. Black dashed lines indicate standard deviation of the ratings. b) The first 5 rows and columns of **V** for Asian (top row), Black (middle row) and White (bottom row) faces using an Asian (left column), Black (middle column) and White (left column) representation. Diagonal elements of **V** (i.e. dot products (dp) of PCs) are on average larger for same race faces than other race faces, indicating that the PCs are better aligned for same race than other race faces.

**Figure 7:**
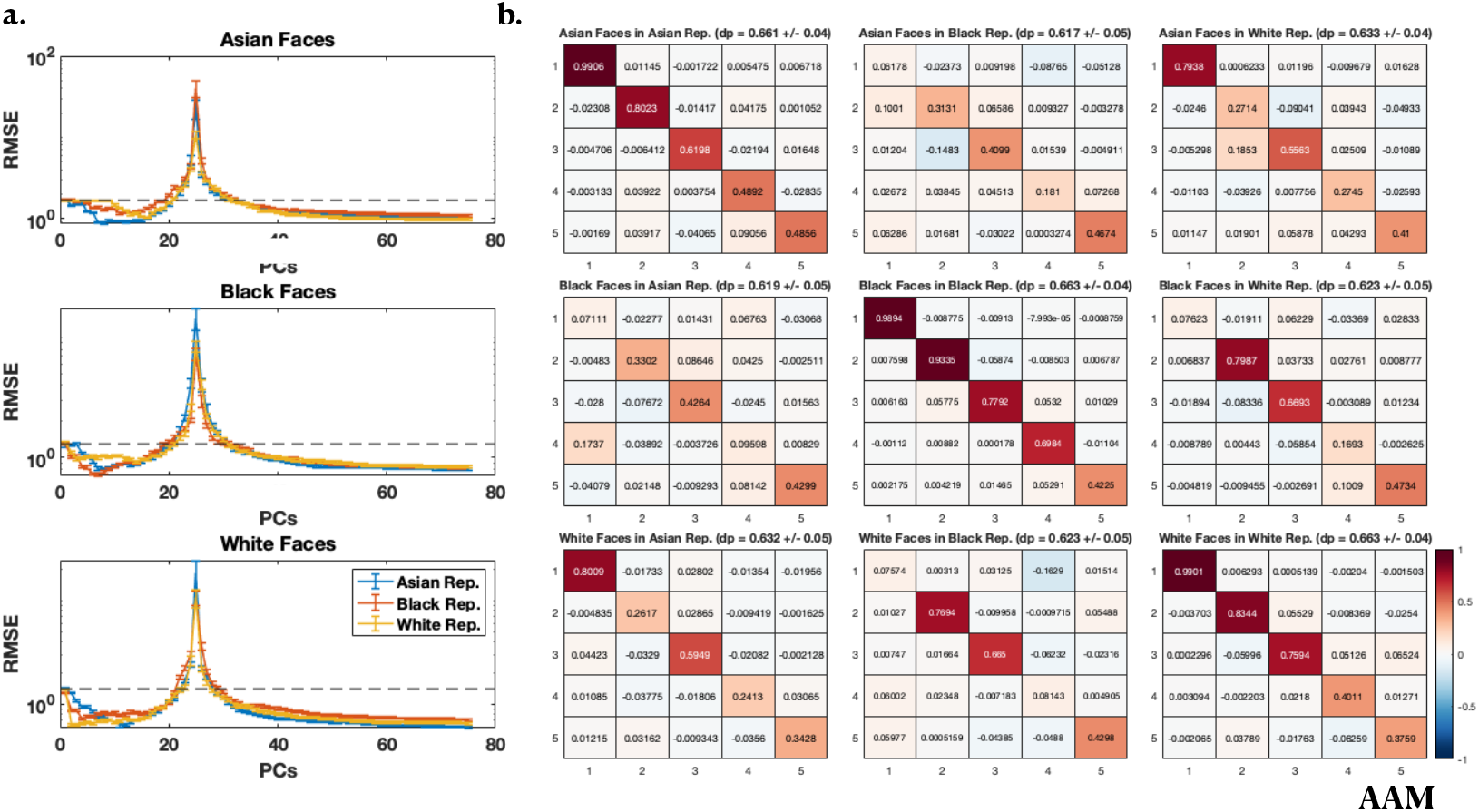
AAM. a) RMSE vs number of parameters for Asian (top row), Black (middle row) and White (bottom row) labeled faces using an Asian (blue), Black (red) and White (yellow) unlabeled face representation for estimating the “feminine” trait from Ma et al. (2015). Performance is better for same race faces than other race faces, indicating that ORE is present in all cases. Black dashed lines indicate standard deviation of the ratings. b) The first 5 rows and columns of **V** for Asian (top row), Black (middle row) and White (bottom row) faces using an Asian (left column), Black (middle column) and White (left column) representation. Diagonal elements of **V** (i.e. dot products (dp) of PCs) are on average larger for same race faces than other race faces, indicating that the PCs are better aligned for same race than other race faces.

